# Spring Model – chromatin modeling tool based on OpenMM

**DOI:** 10.1101/642322

**Authors:** Michal Kadlof, Julia Rozycka, Dariusz Plewczynski

**Author notes:** Email addresses (Michal Kadlof), (Julia Rozycka).

## Abstract

Chromatin structures modelling is a rapidly developing field. Parallel to the enormous growth of available experimental data, there is a growing need of building and visualizing 3D structures of nuclei, chromosomes, chromatin domains, and single loops associated with particular genes locus. Here we present a tool for chromatin domain modeling. It is available as a webserver and standalone python script. Our tool is based on molecular mechanics. It uses OpenMM engine for building models. In this method, the user has to provide contacts and will obtain 3D structure that satisfies these contacts. Additional extra parameters allow controlling fibre stiffness, type of initial structure, resolution. There are also options for structure refinement, and modelling in a spherical container. A user may provide contacts using beads indices, or paste interactions in genome coordinates from BEDPE file. After modelling user is able to download the structure in Protein Data Bank (PDB) file format for further analysis.

We dedicate this tool for all who are interested in chromatin structures. It is suitable for quick visualization of datasets, studying the impact of structural variants (SVs), inspecting the effects of adding and removing particular contacts, measuring features like maximum distances between certain sites (e.g. promotor-enhancer) or local density of chromatin.

## 1. Introduction

Nowadays we observe an enormous increase of interest in the 3D chromatin structure. Novel techniques based on chromosome conformation capture (3C) technology, like Hi-C or Chromatin Interaction Analysis by Paired-End Tag Sequencing (ChIA-PET) provided a great amount of data related to internal contacts, with great resolution, up to ∼1 kbp[1–4]. Since the first data sets appeared, there have been efforts to extract relevant biological information from them, using various bioinformatic and statistical techniques[5]. At the same time, many scientists have tried to determine the exact internal structure of the nucleus for multiple reasons. The most importantly obtaining a model that can explain observable data is confirmation of deep understanding of biological mechanism ruling a structure. Secondly a proper 3D model will be able to predict the effects of SVs and mutation effects, and in future directly connect the structure changes with gene expression levels. Thirdly 3D models are useful for visualization of studied regions..

Multiple modeling techniques have appeared, based on different assumptions, backed up by different theories. One of the earliest modern models of chromatin was based o Grossberg’s[6] crumpled globule, and recapitulated by Mirny in his studies on fractal globules as chromatin models[7]. A completely different approach was applied in [8, 9] where authors employed multidimensional scaling (MDS) on transformed Hi-C maps. The idea behind this technique is that, if we have matrix with distances (similarities) between points, we can distribute these points in 3D space to satisfy these distances. It is like getting a map with positions of cities just from a table of distances between them. However transforming Hi-C matrices into distances is a non-trivial task.

Another group of methods, based on the observation that chromatin loops in Bilateria animals cells are formed in presence of some mediators. One of the most frequently mentioned is CCCTC-binding factor (CTCF) also known as 11-zinc finger protein. In this approach the loop is the primary element of model. In the loop-extrusion-model the chromatin fiber is dynamically extruded through a ring of cohesin and stops after meeting a CTCF motif with the necessary orientation[10]. Another model was developed by group led by Mario Nicodemi called the String-and-Binders-Switch model (SBS), in which chromatin fiber is surrounded by free floating binders that may interact with certain sites on the fiber thus making it stick together[11]. Another Monte Carlo method based directly on ChIA-PET PET clusters is 3D-GNOME, and is implemented as a webserver [12]. Last, but not least, it is worthwhile to mention the Maximum Entropy model, which finds an ensemble of chromosome conformations consistent with a pseudo-Boltzmann distribution for an energy landscape that reproduces the experimentally measured pairwise contact frequencies from Hi-C maps[13].

In this paper we introduce another chromatin conformation modeling tool – Spring Model (SM), a fast, simple to use and powerful tool for visualisation of a fiber with a given set of contacts, in 3D space. We dedicate this tool for everyone who would like to analyse their region-of-interests (ROI) from a structural perspective. It allows one to observe, how a structure will change after adding or removing some interaction sites, simulating effects of structural variants (SVs), and thus observing how changes of distances between enhancer and promoter may affect the expression level of genes. Our tool is based on molecular mechanics (MM) and employs the OpenMM framework[14]. The only required information is a set of contacts and optionally, directionality of CTCF motifs in the sequence. As a result, the user obtains structural data in Protein Data Bank (PDB) file format, which is convenient for further analyses in many popular software packages analyzing biomolecular structures, foe example UCSF Chimera[15]. Our tool is open-source and publicly available for both, on-line usage as a web service, and stand-alone application, that can be downloaded and tuned if so needed, for more advanced usage cases.

Web-service: www.spring-model.cent.uw.edu.pl Stand-alone version: https://bitbucket.org/4dnucleome/md_soft/

## 2. Method description

In SM a polymer is represented as a set of connected beads in 3D space. Their centers are distributed uniformly across the polymer chain i.e. distances between any two consecutive beads are constant and equal in bead diameter. An initial structure must be generated; self-avoiding random walk (SAWR) are a good choice for this task. The other generators of initial structure are also available for the user’s convenience (see section 6). Information about the contacts must then be provided. A contact is defined as a pair of indices, of beads i and j that are supposed to be close to each other in the resulting structure. There are many different ways to extract contacts from raw data, and this part is described more detailed in section 3.

### 2.1 Modeling engine and polymer energy

The identified contacts are added to a system as virtual springs (i.e. harmonic bonds), that are initially stretched. The energy of the system is then minimized, springs shorten and fiber folds into a structure that satisfies the chosen contacts. A schematic of the concept is depicted by fig. 1.

**Figure 1:**
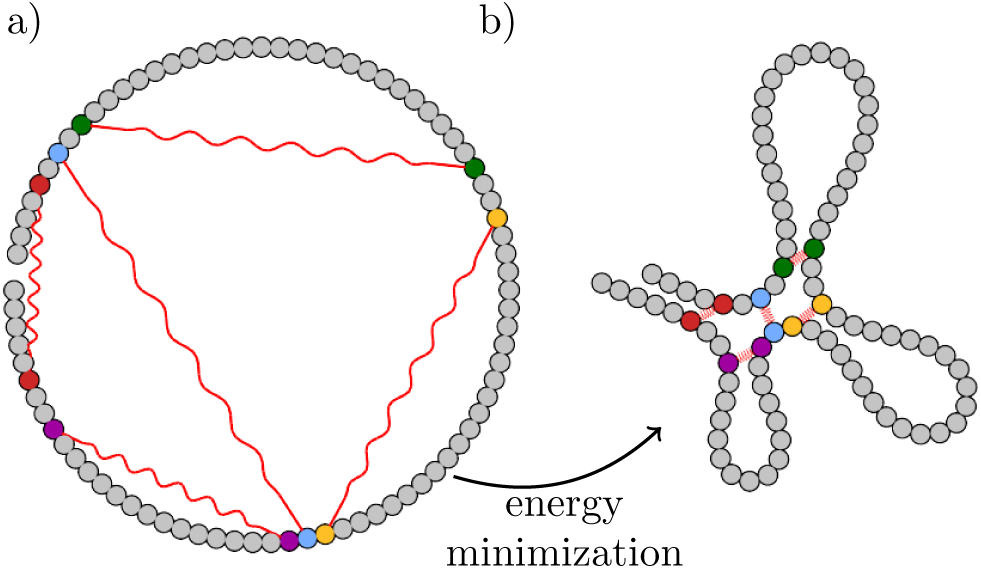
a) Initial structure. The initial structure depicted as a circle for clarity, is in most cases a self-avoiding random walk. Virtual springs are depicted as red sinusoidal lines. Connected beads share the same color. b) structure after energy minimisation.

Energy in spring model is defined as a sum of following terms, described in detail below:

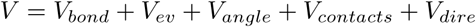

*V* is the hipersurface of potential energy. It is a function of beads’ positions in the system. Minimizing this function will lead us to find a local minimum in such surface and thus a structure that satisfies the given constraints.

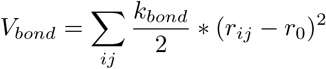

*V*_*bond*_ is a potential between consecutive beads. Force constant *k*_*bond*_ is arbitrarily chosen to be high enough so as to treat the length between beads as constant. The equilibrium constant *r*_0_ (bond length) is equal to the bead diameter *σ* (by default 1) and *r*_*ij*_ is at the same time the distance between beads i and j.

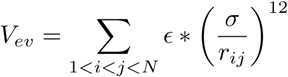

*V*_*ev*_ is responsible for excluded volume (EV). In SM we assumed that chromatin fiber can not cross itself and each bead represents a physical amount of chromatin, possessing volume which cannot overlap with other beads. To achieve this we introduce the EV term defined as the repulsive part of Lenard-Jones potential. *ϵ* is a scaling factor chosen arbitrarily, and *σ* is the bead radius.

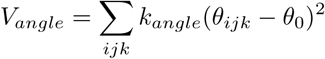

*V*_*angle*_ is a harmonic 3-body angle potential that gives the fiber it’s stiffness. *k*_*angle*_ is the force constant, *θ*_*ijk*_ the flat angle between consecutive beads i,j,k, and *θ*_0_ the equilibrium constant set to *pi* rad. These terms make the fiber straighten and are overcome by the contacts term. *k*_*angle*_ is a parameter that strongly depends on model resolution (at lower resolutions it can be completely omitted since we can assume that unconstrained regions of regions of chromatin behave like a pure random walk. It can also be used to make loops in the model more clearly visible

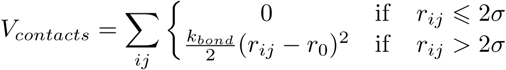

*V*_*contacts*_ is the contact’s potential. It’s very similar to *V*_*bond*_, however since it has a flat dip, it allows a little more freedom for bead interactions.

*V*_*dire*_, the directionality potential that allows additional loop shape optimization. This is a 4-body, dihedral angle, harmonic potential between interacting beads and their closest neighbors. It is described in more details in section 2.2.

### 2.2. CTCF motifs directionality and loop types

The CTCF protein recognizes appropriate motifs on DNA sequence. The directionality of these motifs is essential for forming chromatin loops, and higher level domains [16, 17]. As it was stated in [18] in the cell nucleus, there is a possibility for occurrence of two types of loops - *coiled* (C-loops), (fig. 2) and *hairpin* (H-loops). The type of loop is dependent on CTCF motifs directionality. If a motif is convergent (><) then H-loops are expected to form, whereas divergent loops do not form at all (<>). In case of motifs situated as tandem left or tandem right(<<, >>) C-loops are expected. To reflect this behavior, our methodology implements additional an force, which can be applied to bend the fiber accordingly for further loop shape optimization.

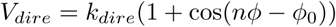

**Figure 2:**
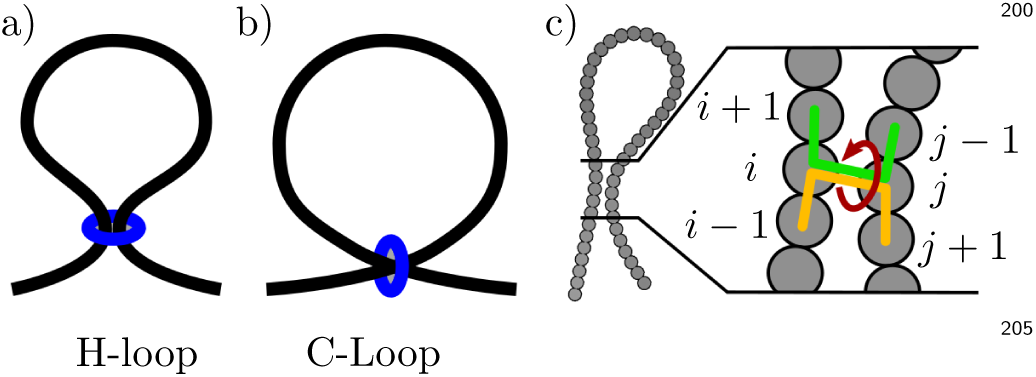
Loop types: a) Hairpin loop, b) Coiled loop. Blue rings represent binding sites. c) dihedral angles responsible for forming different loops types. Two separate terms engaging different beads are marked with a green-yellow line. Example for H-loop is depicted.

This extra term can be provided per interaction and is implemented as *periodic harmonic dihedral angle force* defined between consecutive beads *i* − 1, *i, j, j* + 1 and *i* +^195^ 1, *i, j, j* − 1 with an equilibrium constant equal to 0 rad for C-loops, and *π* rad for H-loops, and force constant is equal to 50 kJ/mol.

## 3. From experimental data to set of interaction

The Spring Model was designed as a tool for building a 3D conformation that satisfy a given set of constraints. It should be used as a tool for visualization certain domain in single cell rather than population based average model as it take place in [12]. This require careful data selection and sometimes filtering. Applying to much interactions on short region of chromatin will lead to collapsing model (see fig 3). Contacts may be derived from great variety types of experiment, e.g, ChIA-PET, Hi-C, single-cell Hi-C or even ChIP-Seq processed by machine learning techniques [19]. There are several ways to extract interactions from raw data and exact method is strongly dependent on data and expected results. In following paragraphs different types of data will be characterized in the light of their usage in modelling process.

**Figure 3:**
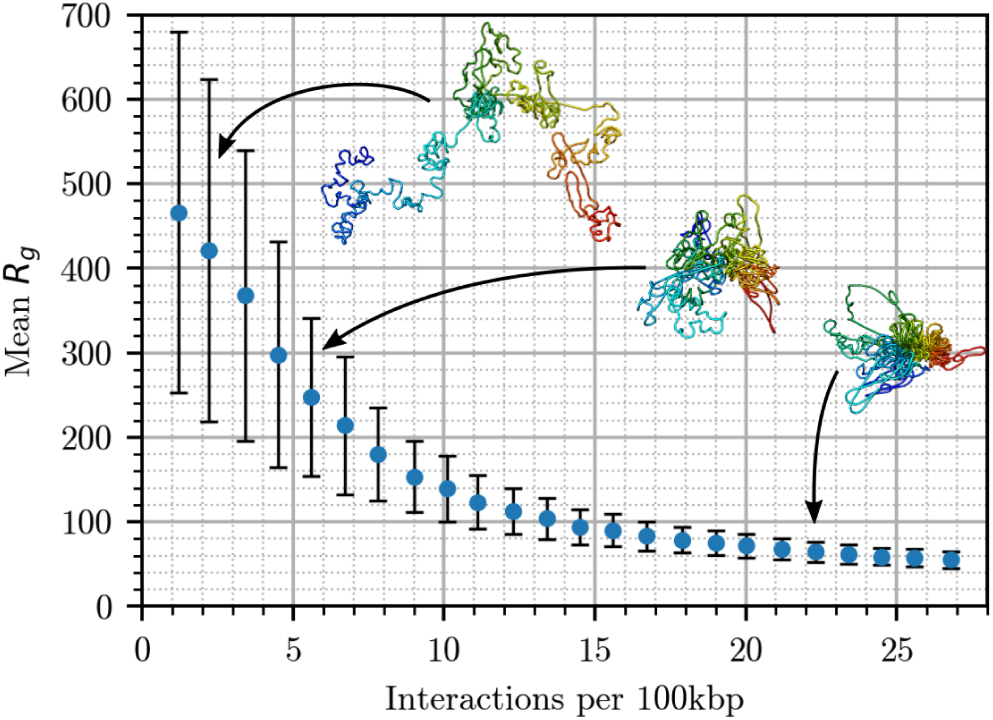
Radius of gyration as a function of number of interactions per 100 kbp. High *R_g_* are indicative of a poorly folded structure, and a low *R_g_* indicates a fully collapsed structure. Error bars indicate the standard deviation. There is no clear phase transition point. Representative structures are depicted for three clusters.

### 3.1. Hi-C

Hi-C are population based experiment in which one observes frequencies of all possible contacts (even accidental). It is impossible that all of them exist in the same time in single cell, thus some sampling from heat map is required. The simplest approach is just cutting off an interaction fre-quency heat map at an arbitrary level and taking what’s left – the strongest (i.e. most confident) interactions. Fur-ther studies on decomposition Hi-C heatmaps into separate cell is required.

### 3.2. scHi-C

Despite the development of modern experimental methods of studying the structure of the genome, several questions (important especially from a modeling point of view) remain unanswered. One of the most important is: how many contacts observed in Hi-C/ChIA-PET, or other similar experiments are actually realized in a single nucleus? The main challenge is that the aforementioned techniques give averages over millions of cells. Although there are some approaches to identify contacts in single cells [20, 21], we are still unsure how many contacts remain hidden. Single Cell Hi-C seems to be perfect technique for modelling purposes. However the main drawback of this method is that we don’t know how much contacts that really occurs we don’t see. According to our initial proof-of-the-concept tests based on data published by [20] our conclusions where that this set is not big enough to obtain well folded structures (detailed data not published).

### 3.3. ChIA-PET

ChIA-PET is another population based technique which shows contacts mediated by particular protein. It returns in theory a subset of contacts visible in Hi-C. It also poses much better resolution, and shows direct interactions between particular sites in chromatin. In experiments with shallow sequencing and low number of loops all interactions may be used directly. In experiments with deep sequencing, and large number of loops some sampling is required. We suggest randomly selecting the loops with probability proportional to PET-Count. This also recommended way for building ensemble of solutions if needed. Yet another approach was proposed in [22].

### 3.4. Impact of number of interactions

What we do know is that some contacts occur more often than others and they cannot all occur at the same time. Thus there ought to exist an average amount of chromatin per interaction, or in language of model coordinates – number of beads per interaction (BPI). What is more, the values from this parameter are closely related to the chromatin state. The active states of chromatin are associated with higher loop density [18].

To explore impact of BPI on structure, we measured the mean radius of gyration *R*_*g*_, over ensembles of 1 000 generated models for our ROI (chr6:32047146-33250152), where 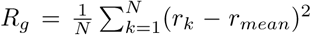, and *N* is number of beads, *r*_*k*_ is a vector indicating positions of *k*th bead, and *r*_*mean*_ is the geometrical (which in this case implies gravitational) center of a system. For our ROI we have selected CCDs with length in range (1-2 Mbp) that have the highest percentage of its length covered by regulatory elements. The regulatory elements were defined based on Combined ENCODE Segmentation[23]. We have included into this category following classes of regions: CTCF enriched element (CTCF), Predicted enhancer (E), Predicted promoter flanking region (PF), Predicted promoter region including TSS (TSS), Predicted weak enhancer or open chromatin cis-regulatory element (WE). For the set of experimentally resolved interactions (ChIA-PET for GM12878 cell line mapped onto hg38 genome – data not published), we were generating ensembles of models. Each model was starting from a different initial structure (SARW), and in each ensemble we chose a different number of contacts included in modeling. Contacts were randomly chosen with probability proportional to PET Count from ChIA-PET experiment. The results are depicted on fig. 3.

We did not observe any clear threshold that could be associated with phase transition from a “folded” to an “unfolded” state. However it has not escaped our notice that the specific threshold of the number of used interactions induces almost the same level of chromatin compaction. We conclude that no more than 15 interactions per 100 000 base pairs take part in forming chromatin structures in particular cells. Adding more contacts does not significantly change the level of compaction.

## 4. Modeling example

As an example of a use case of our modeling tool we chose TAL1 locus, a proto-oncogene activated by disruption one of CTCF sites between TAL1 and a nearby enchancer located on the CMPK1 gene [24]. The authors identified 15 ChIA-PET interactions in this region, and after ruling out one of anchors only 9 interactions remain. We are aware that the number of interactions in ChIA-PET experiments depends on sequencing depth. However, due to the small amount of loops we decided to use all of them in modeling just for visualisation purposes. We built both: a wild type model, and a model after deletion of CTCF using CRISPR. The results are depicted in the fig. 4.

**Figure 4:**
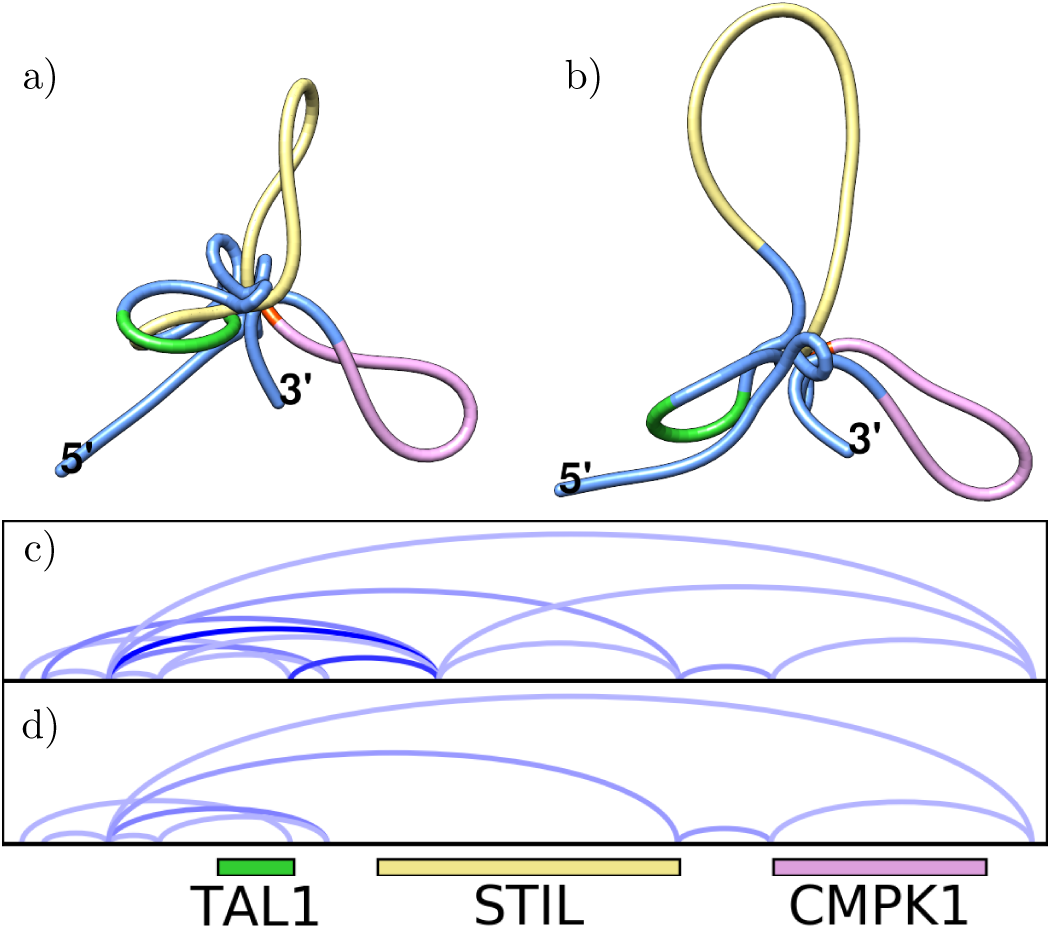
Models of Tal1 locus for a) wild type, b) CTCF deleted with CRISPR. c) loop diagram diagram for WT, and d) ΔCTCF. Loop color saturation corresponds to PET-Count. Colors on models corresponds to colors of genes marked below arc diagrams. Deletion of CTCF binding site leads to desappear of 6 contacts which result with less compacted structure and more flexible loops.

It’s clearly visible that even if a certain anchor is deleted and some loops disappear, other can still act as an insulator between the TAL1 promoter and enhancer near CMPK1, since they still remain on distinct loops. More effort needs to be put to explore the methods of loop sampling from ChIA-PET to make this data suitable for modeling purposes.

## 5. Validation

Even in today’s current chromatin modeling field, there is still big deficiency of structure validating methods. It is hard even to define the criteria to be used in assessment of the reliability of a model. This deficiency is caused by lack of methods for direct study of chromatin structure (analogous to X-Ray crystallography in the case of proteins). Although there is considerable progress in the field of direct 3D imaging[25, 26], building image-driven models consistent with genomic data remains an open problem.

We access the accuracy of our modelling procedure based on the ability of proposed models to recreate the large - scale features observed for the specific cell line. To check the consistency of our approach with ChIA-PET experiment data, we generated 10 000 models starting from SARW and randomly choosing interactions with probability proportional to the PET-Count, ROI being chr7:141246953-142936866 (hg38). Then for each structure we calculate the binary contact map and sum them up. The resulting matrix bears many similarities to the original heatmap with the chessboard-like structure and enhanced signal along the loops boundaries. (fig. 5) with contact frequencies, which we assume meet the user’s expectations. We call this procedure simulated ChIA-PET. Similarly to the real experiment, only an assembly of a large enough number of models can reproduce population characteristics of the data.

**Figure 5:**
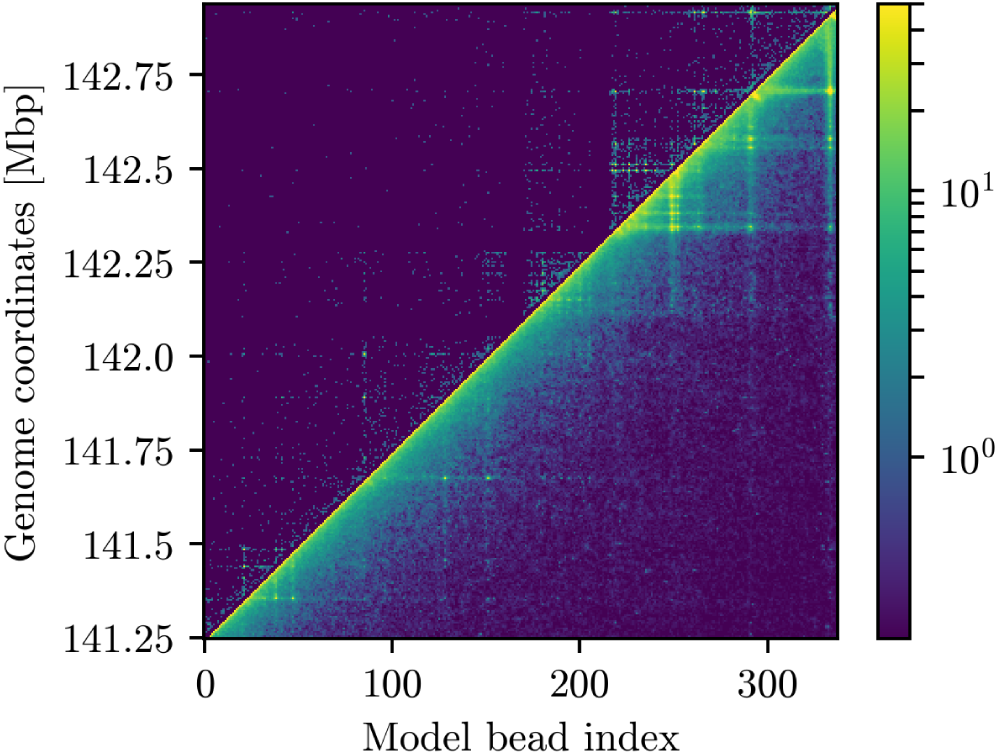
Upper half of the heatmap shows raw ChIA-PET data that was used to model the region (chr7:141246953-142936866). Lower half depicts population-scale contact matrix obtained from summing binary contact matrices from 10 000 structures built with the Spring Model. Both matrices were normalised to convey the relative frequency of the contacts; the scale is logarithmic with base 10, dark purple being little to no contacts and yellow representing the other side of the spectrum.

In [19] we showed that there is a possibility of predicting ChIA-PET interactions from data from cheaper experiments like ChIP-Seq, and in many cases, the interaction pattern is analogous to similarities between 3D structures. We also found out that in some cases it is easy to radically change the structure by simply removing one key long range contact; whilst in other some contacts seems to be redundant because even if they are missing other neighborhood interactions keep structure folded.

## 6. Webservice

For user convenience our methodology was implemented as a web service available at http://spring-model.cent.uw.edu.pl. It was designed for being simple and intuitive. The landing page is a form for submitting new tasks - see Fig.6. The user has a wide variety of options that allow him or her to tailor his model according to his needs. First of all a user is able to name accordingly his task which help him easy identification later. Then the user is able to choose one of four initial structure types: (almost) straight line, circle, self-avoiding random walk and baseball. Then there are options for controlling stiffness of the fiber and size of the spherical container if it is needed. In rare cases depending on provided data and initial structure, simple molecular mechanics may have trouble with convergence to a reasonable structure, and some artifacts, or not completely folded structure may occurs. To over-pass this phenomenon, there is an option for running modeling with refinement, which means that after the first act of energy minimization there will be another 5 more cycles of running short (1000 steps) molecular dynamics (with time step Δ*t* = 20 fs) in a temperature of 300 K and energy minimization with convergence condition equal 10 kJ/mol, and in the end yet another energy minimization run with convergence condition equal 1 kJ/mol. This usually allows to achieve lower energies of the final structure however, it costs more computational power and may extend computation time from seconds to minutes depending on system.

**Figure 6:**
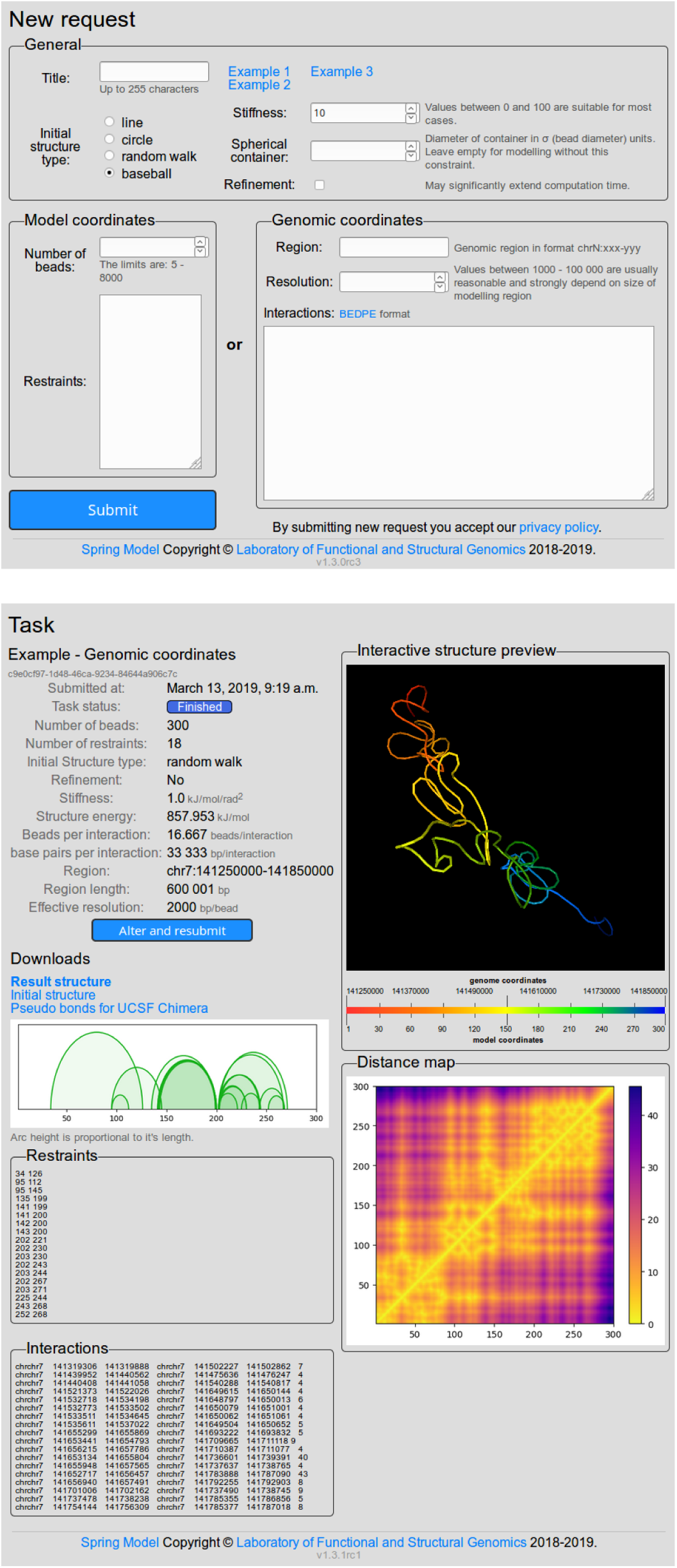
Web service interface. New request form, and example of modeling results.

In the second part of the interface, user’s have an option to chose how he is going to express restraints. There are two options. One may directly provide indices of interacting beads, or use interactions expressed in genome coordinates saved in BEDPE file format. In first option a user can additionally provide number of beads. If this parameter will be skipped the number of beads will be equal to the highest index of bead in the list of restraints. After indices, the user may specify expected type of loop (as described in section 2.2). Using symbols of CTCF motif directionality >>, ><. <>, <<), or upper case letter (C for coiled loop, or H for hairpin loop). As another option, user have to additionally provide information about resolution i.e. amount of chromatin. Analogically fiber length can be controlled using a region of field, or it will be deduced from coordinates. Loop types can be specified as 8^th^ column of a BEDPE file. In this case, the midpoints between starts and ends of anchors will be used for converting input data to restraints.

After few seconds to minutes, users can inspect the results on a results view presented on Fig.6. There is a multiple useful information about structure, like basic statistics – number of beads, restraints, number of beads per interaction, region length and resolution, type of initial structure, stiffness, size of spherical container, and energy of the final structure. There is also the visual representation of interactions as an arc diagram and interactive preview of structure in 3D, that allows to rotate, zoom in and zoom out the structure. If user is not satisfied with obtained model or would like to experiment with different parameters there is convenient button that redirects back to the new task form that is already populated with data from a task. For more advanced structural analysis user can download the structure in PDB file format which is widely supported by most of modern bimolecular viewer and analysis software, e.g. UCSF Chimera [15]. Additionally the user may download files with restraints saved as a pseudo bonds file format that can be rendered in UCSF Chimera. This allows for the clear examination of positions of interactions in the structure. As the final option, users may refer to provided lists of restraints or couple them with genomic interactions. Users may also visually inspect distance maps of the resultant structures.

## Acknowledgment

This work has been supported by Polish National Science Centre (2014/15/B/ST6/05082), Foundation for Polish Science (TEAM to DP). The work was co-supported by grant 1U54DK107967-01 “Nucleome Positioning System for Spatiotemporal Genome Organization and Regulation? within 4DNucleome NIH program, and by European Commission as European Cooperation in Science and Technology COST actions: CA18127 “International Nucleome Consortium” (INC), and CA16212 “Impact of Nuclear Domains On Gene Expression and Plant Traits”. Authors would like to express our special thanks of gratitude to our colleagues from the laboratory who have critically examined the first version of this manuscript, and did necessary language corrections.

